# PTY 2 As DPP-IV Inhibitor Prevents Intestinal Cell’s Apoptosis

**DOI:** 10.1101/670430

**Authors:** Shivani Srivastava, Harsh Pandey, Surya K Singh, Yamini Bhusan Tripathi

**Affiliations:** Department of Medicinal Chemistry, Institute of Medical Sciences, Banaras Hindu University, India; Department of Endocrinology and Metabolism, Institute of Medical Sciences, Banaras Hindu University, India

**Keywords:** *Pueraria tuberosa*, Diabetic Complications, DPP-IV, Streptozotocin, Apoptosis

## Abstract

Enhanced DPP-IV expression is found to get enhanced in various intestinal diseases. PTY-2 has already been known to have DPP-IV inhibitory potential. This inhibition has not yet been studied at mRNA level. Increased incretin secretion due to DPP-IV inhibition could lead to suppression of stress & intestinal cells apoptosis. Through histological analysis, we have found morphological damage of intestine after STZ injection (65 mg/kg bw) to male Charles foster rats. mRNA expressions were analyzed by PCR and apoptosis of cells was checked through tunnel assay & Bcl 2 expression. The number and length of villi get reduced in STZ induced diabetic control, but these damages have been getting reversed after the PTY-2 treatment for 10 days. The expression of SOD was found to get reduced while that of DPP-IV was enhanced in diabetic control group along with significant intestinal cell apoptosis. PTY-2 treatment reduces the stress by upregulating the expression of SOD and through downregulation of DPP-IV mRNA expression. These recoveries by PTY-2 leads to suppression of intestinal cells apoptosis. These short studies explain the protective action of PTY-2 against STZ induced intestinal damage. Hence, PTY-2 could be taken as an herbal treatment against intestinal disorders in case of chronic diabetes.

## 1. BACKGROUND

Since 1963, STZ has been used to induce diabetic experimental models. The STZ enters via GLUT 2 transporter and harms β cells through DNA methylation and by acting as a nitric oxide donor. In addition to pancreatic β cells, the GLUT 2 transporters are also responsible for STZ uptake into the epitheliocytes of the intestinal mucosa, cells of renal tubules and hepatocytes. It means that, STZ is toxic to other organs on those cells which express the GLUT 2 transporter(Akinola et al., 2009). Several herbals has been studied to improve the intestinal deformities against STZ induced diabetes like Azadirachta indica(Akinola et al., 2009), Green tea extract and ginseng root(Karaca et al., 2011).

According to the previous study, the differentiation-dependent expression of DPP-IV in crypt-villus axis of rat jejunum is primarily controlled at the level of mRNA(Darmoul et al., 1991). Promotion and formation of intestinal ulcers are also known to be inhibited by DPP-IV inhibition(Inoue et al., 2014). GLP-2, a hormone responsible for intestinal growth is usually found to be inhibited by DPP-IV in rats. The DPP-IV inhibition by valine-pyrrolidide (VP), reduces the GLP-2 degradation and enhances its intestinotrophic effect (Hartmann et al., 2000).

We have already reported the DPP-IV inhibitory role of PTY 2 in blood and intestinal homogenate(Srivastava et al., 2015, 2017, 2018b). Here, we have further studied the mRNA expression of DPP-IV in the intestinal duodenum and used this property in designing our study focused to STZ induced intestinal damage. *Pueraria tuberosa* have positive role as antioxidant(Pandey and Tripathi, 2010) & anti-inflammatory(Pandey et al., 2013) agent. It is highly effective in the treatment of diabetes(Srivastava et al., 2015, 2017, 2018a), nephropathy(Tripathi et al., 2017), anti-hypertension(Verma et al., 2012), anxiolytic(Pramanik et al., 2010) etc. Steroid, glycoside, triterpenoid, flavanoid, carbohydrate, tannin, protein, alkaloids and amino acids are the main constituents of *Pueraria tuberosa* like puerarin 4’,6’-diacetate, tuberosin, daidzin, genistein, puerarin, puetuberosanol, puerarone & tuberostan(Asthana et al., 2015; Srivastava et al., 2015; Tripathi and Kohli, 2013). We have hypothesized that PTY 2 treatment could be responsible for the improvement of intestinal morphology against STZ induced damage.

## 2. MATERIALS AND METHODS

### 2.1 MATERIALS

For RT-PCR, Trizol (Himedia, Pvt. Ltd,Kolkata, India), cDNA Kit (Fermentas), and Taq-polymerase (GenaxyScientific Pvt.Ltd) were used.

### 2.2 SAMPLE PREPARATION

*Pueraria tuberosa* was purchased from Ayurvedic Pharmacy, Banaras Hindu University. Its authenticity has already been ascertained in our previous research (Pandey and Tripathi, 2010). 30 g of tuber powder were extracted with 8 volumes of distilled water. When the volume reduced to one-fourth, the obtained extract then filtered. The total yield obtained was 30% (Srivastava et al., 2018a).

### 2.3 ANIMALS DESIGN

After acclimatization, charles foster male rats of the same age group and a weight range of 120-130 g were injected STZ (65 mg/kg bw), prepared in fresh and chilled citrate buffer (pH 4.5), after fasting for 8 hrs. With the use of glucometer strips (Dr. Morepen), the diabetic condition were checked on the 5th day. The rats with blood glucose level above 200 mg/dL were considered diabetic and kept for 60 days in order to induce chronic diabetes. On 61th day, the rats were divided into three groups (n=6): Group-1 (STZ untreated rats, i.e., age-matched normal control), Group-2 (diabetic control), and Group-3 (PTY-2 at 50 mg/100 g bw treatment for 10 days to diabetic rats). After 10 days, the rats were sacrificed and intestine samples were isolated. Each intestine sample was cut into two parts; one for histology and IHC (preserved in 10% formaldehyde) and the other was first crushed in liquid nitrogen and then stored in −80°C freezer for molecular study(Srivastava et al., 2018a).

### 2.4 HEMATOXYLIN-EOSIN (H & E) STANING

Intestinal tissues fixed with formalin were embedded in paraffin wax. Using Leica RM2125 RT rotator microtome (Leica Biosystems Nussloch GmbH, Nussloch, Germany), the tissues were cut into 4 µm thick sections. Each sections, were latter stained with hematoxylin-eosin (HE) and then imaged & observed using Nikon microscope (Eclipse 50i, loaded with imaging software-NIS Elements Basic research).

### 2.5 TUNNEL ASSAY

Apoptosis assay was done using “TACS^®^ 2 TdT Fluorescein Kit - Trevigen”.

### 2.6 IMMUNO-HISTOCHEMICAL STAINING

Dewaxed the intestinal tissues using xylene for 10 min and through 90 %, 70 % alcohol and water, tissues were then rehydrated serially in each for 5 min. Then after, each slides get dipped in citrate buffer and proceded for antigen retrieval by using EZ Retrieval System V.3 (Bio Genex). Sections were washed two times with PBS for 10 min each, and then blocked with 0.1% Triton X-100, 0.1% BSA, 10% FCS, 0.1% sodium deoxycholate and 0.02% Thiomersal (anti-fungal agent) in 1X PBS for 2 h at room temperature (RT). All sections were then incubated overnight with the primary antibodies at 4°C and then washed with PBST (0.1% triton X in 1XPBS) thrice for 10 min each, followed by incubating each sections with anti-rabbit-AF 546 (Red) (Invitrogen, USA) secondary antibodies at RT for 2 h. Again washed thrice in PBST for 10 min each. Finally, counterstained with DAPI (1 µg/ml DAPI in 1XPBS) and mounted on DABCO. Using Zeiss LSM510 Meta confocal microscope all the slides were examined. Zen Black (2012) software was used for image analysis.

### 2.7 REVERSE TRANSCRIPTION-POLYMERASE CHAIN REACTION (RT-PCR)

Using trizol, we have homogenized 50 mg of intestinal tissue. With random hexamers and superscript II RNase H-reverse transcriptase (RT), 5 μg of total RNA was reverse-transcribed. For DPP-IV, 2µl c-DNA, 0.2 mmol/L dNTPs, 1.5 mmol/L MgCl_2_, 0.5 μmol/L of each primer, 2.5µl 10X PCR buffer and 1U Taq DNA polymerase were used. For Glyceraldehyde 3-phosphate dehydrogenase (GAPDH), 0.1 μmol/L of each primer was used. Expressions optical density were determined and presented as ratio against GAPDH with the help of alpha imager (Bio-Rad). All the expressions were checked in triplicate (Table 1).

**Table 1.**
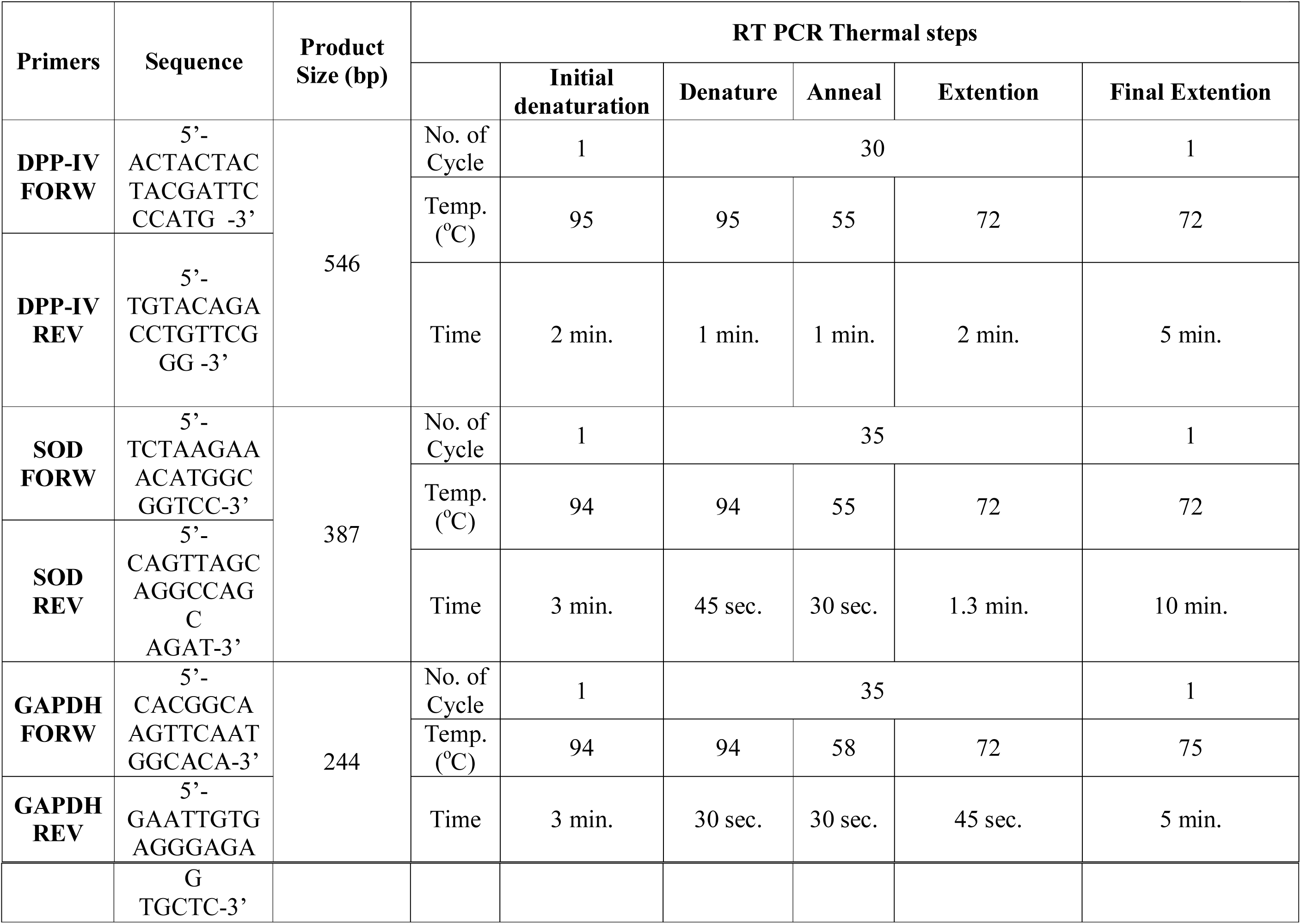
Details of PCR primer sequences, product size and thermal steps for expressions of SOD, DPP-IV and GAPDH.

## 3. RESULTS

### 3.1 HISTOLOGICAL EXAMINATION

STZ treatment reduced the overall length and number of intestinal villi. The PTY 2 significantly prevented/ reversed the STZ induced intestinal changes in 10 days of treatment. This treatment with PTY 2 could be enhancing the surface area of intestine, leading to enhanced absorption of nutrients and minerals (Figure 1).

**Fig. 1.**
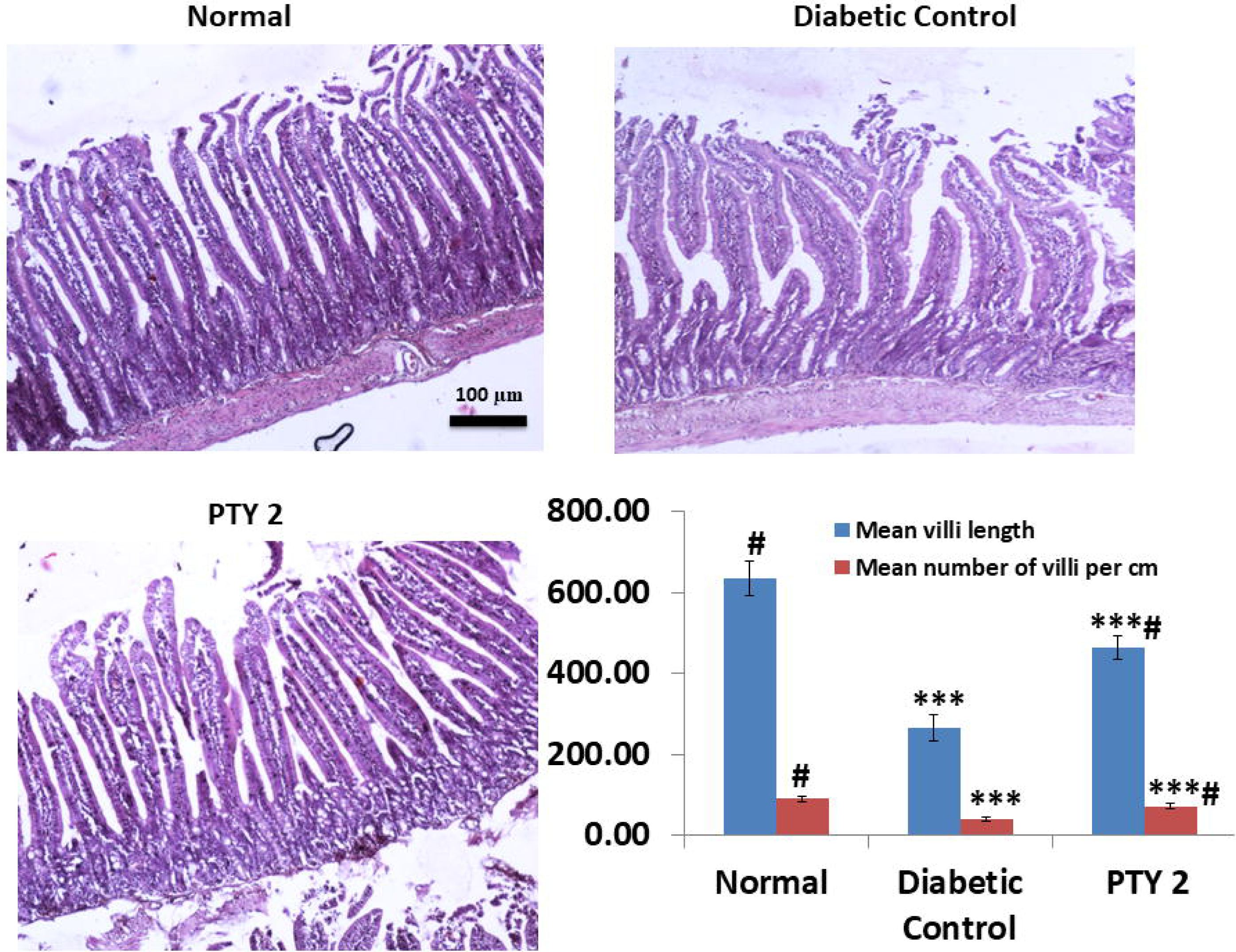
H & E staining to measure the morphological changes of intestinal villi. PTY-2 treatment showed a recovery of STZ-induced villi damage. The image was taken at 20X magnification. Scale bar 100 μm.

### 3.2 APOPTOSIS

Tunnel assay is aimed to assess the degree of apoptosis in cells. The sections of STZ control groups showed significant damage accompanied with cell apoptosis. However, the PTY 2 treatment for 10 days, significantly prevented this change (Figure 2).

**Fig. 2.**
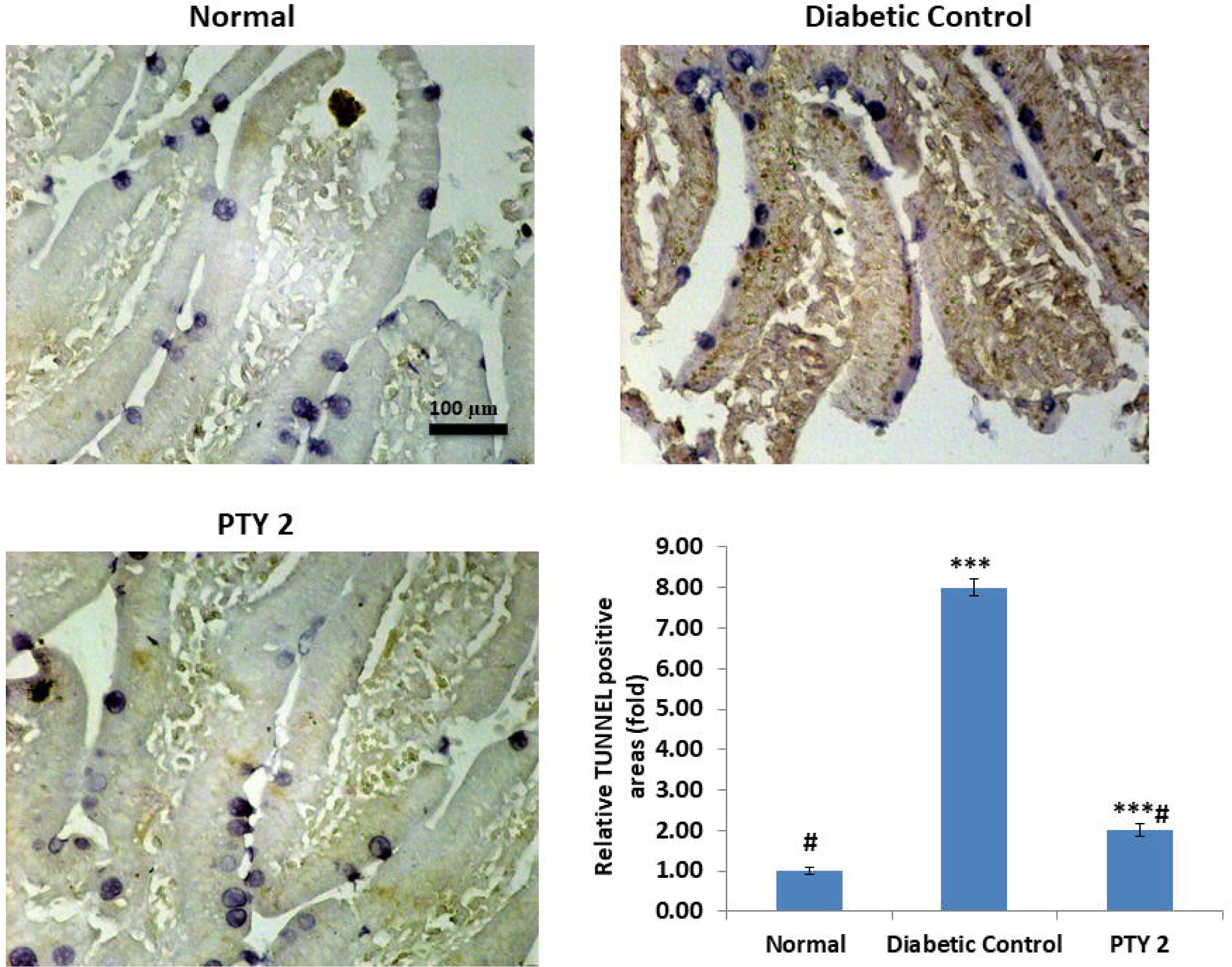
Tunnel assay analysis showed the effect of PTY-2 on intestinal cells apoptosis. PTY 2 recovers the apoptosis induced by STZ. The image was taken at 40X magnification. Scale bar 100 μm. The brownish color indicate the TUNEL-positive area.

Immunohistochemistry analysis also proved the antiapoptotic effect of PTY 2 on intestinal cells. It significantly reversed the downregulation of Bcl 2 expression induced by STZ (Figure 3).

**Fig. 3.**
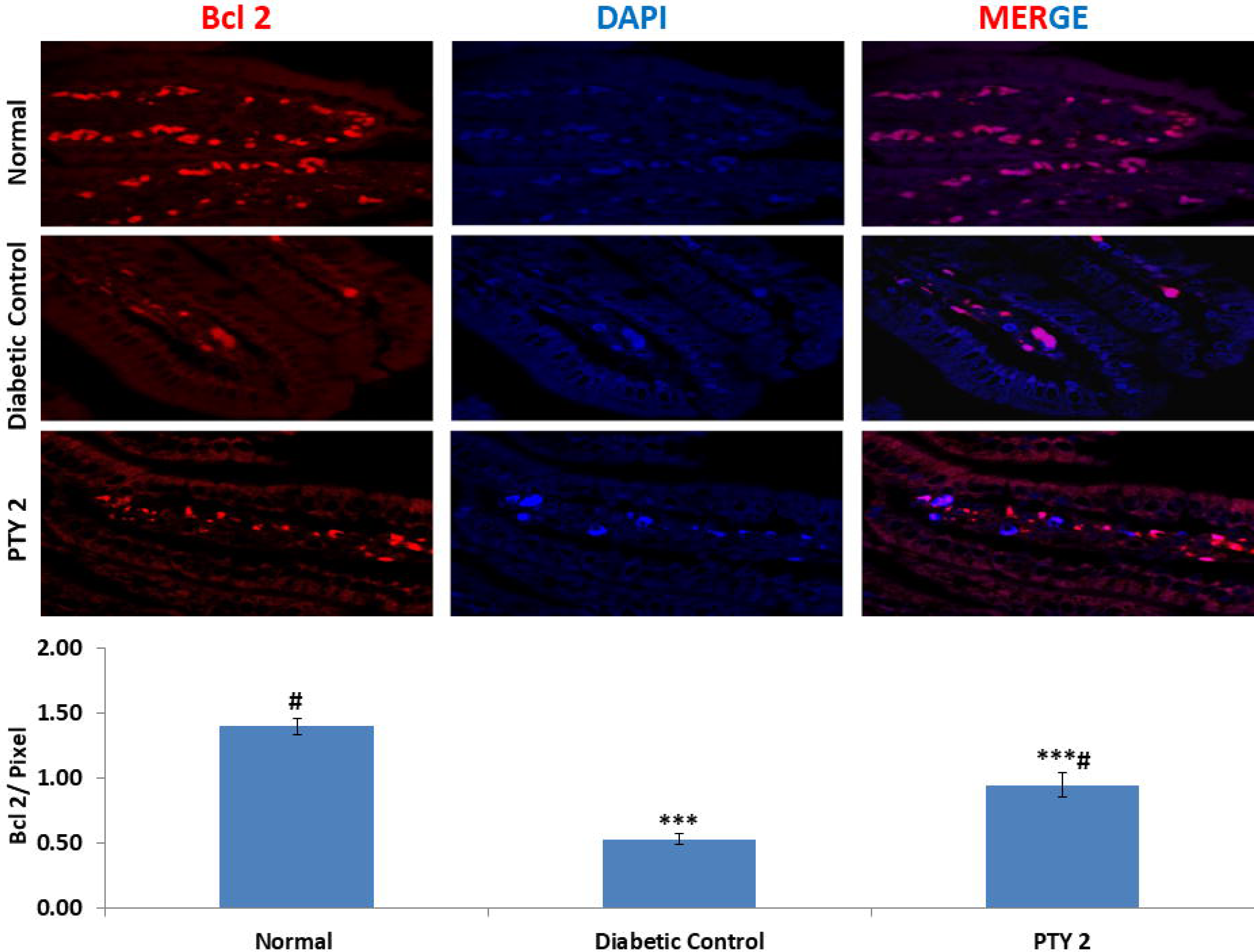
Immunohistochemistry analysis showed the effect of PTY-2 on the expression of Bcl 2 in the intestine of normal, diabetic control, and PTY-2-treated rats. The expression was merged with DAPI (blue). In comparison to diabetic control, PTY-2 up regulated the expression of Bcl 2. The image was taken at 63X magnification. Scale bar was 20 μm. The intensity was measured in pixel values. Each value represent the mean ± SD (n=6); *** *P* < 0.05, compared with normal, # *P* < 0.05, compared with diabetic control.

### 3.3 mRNA EXPRESSIONS

PTY 2 recovers the STZ induced stress, as a result the expression of SOD was significantly enhanced. In the other hand, we have also found the reduced mRNA expression of stress marker DPP-IV in PTY 2 treated group. This clearly shows the potential of PTY 2 against STZ induced intestinal damage (Figure 4).

**Fig. 4.**
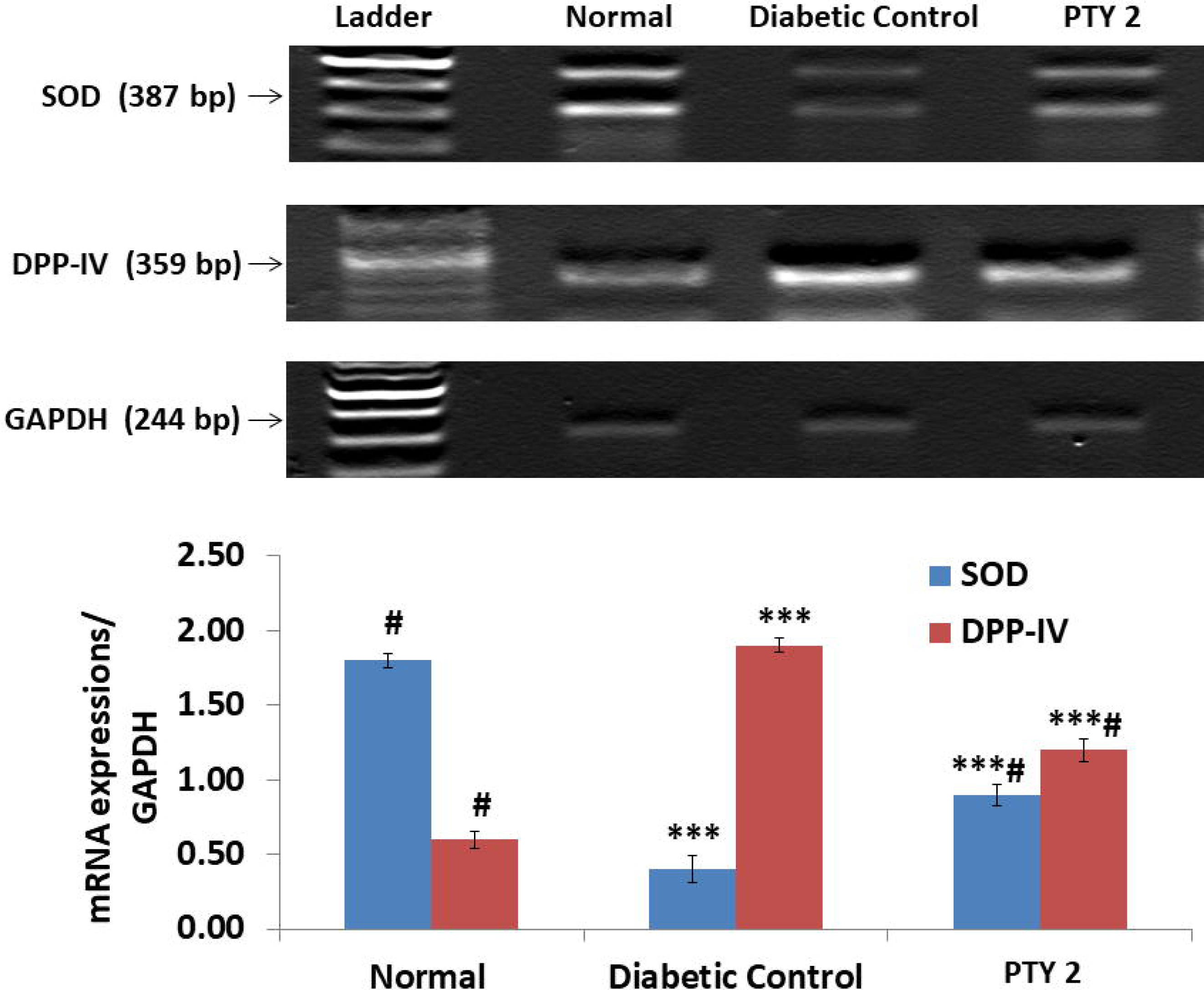
mRNA expression of SOD and DPP-IV to investigate the effect of PTY-2 on the intestinal tissues of normal, diabetic control and PTY-2 treated rats. Each value represent the mean ± SD (n=6); *** *P* < 0.05, compared with normal, # *P* < 0.05, compared with diabetic control.

## 4. DISCUSSIONS

During diabetes, the morphologies and functions of small intestine gets highly altered. Some of this alterations is caused by the oxidative stress raised during the diabetic complications, studied in STZ induced diabetic rats(Bhor et al., 2004).Increase in the intestinal DPP-IV activity is asoociated with diabetes development(Yang et al., 2007). In our case, the enhanced expression of DPP-IV and deformities of intestinal morphology by STZ clearly indicates its high uptake by intestinal mucosa as discussed above. PTY 2 recovers this damages by upregulating the antioxidant enzyme SOD and downregulating DPP-IV mRNA expressions (Figure 4). Thus, reduced stress leads to the significant increase in the number and length of villi as compared to STZ treated group (Figure 1). The apoptosis assay and Bcl 2 protein expression also showed the antiapoptotic and protective effect on intestinal cells by PTY 2 (Figure 2 &3).

Glucagon like peptides plays an important role towards gut adaptation. After their synthesis, they get released into the intestine from the enteroendocrine cells. GLP-1 efficiently assimilates nutrients while GLP 2 act as a regulator of energy absorption, mucosal integrity & permeability(Drucker, 2002). Both GLP-1 and GLP 2 proven to play the essential regenerative and healing role against intestinal injury in mice(Hytting-Andreasen et al., 2018). Through the mechanism involving Fgf 7, GLP-1R improves both small and large bowel growth(Koehler et al., 2015).

In our previous works, we have already proved PTY 2 as incretin therapeutic agent. It significantly inhibits DPP-IV and enhances the levels of GLP-1 and GIP. In addition to DPP-IV inhibition, it also act as incretins receptor agonist(Srivastava et al., 2015, 2017, 2018a, 2018b). Thus, as DPP-IV inhibitor, PTY 2 must also enhances the intestinotrophic effect of GLP 2, in addition to GLP-1 and its receptor.

In sprague dawley diabetic rat model, STZ altererd the microbiota compostions and decreased the microbial diversity with time(Patterson et al., 2015). PTY 2 improves the villi count and length, thus enhances the surface area in order to assimilate more nutrients from diet and could also provide the maximum space for colonization of positive bacteria useful for intestinal health. A detail study is needed to reveal the role of PTY 2 and its individual active constituents in future at both molecular and microbial level against diabetic complications.

## 5. CONCLUSION

PTY 2 recovers the STZ induced stress, improves the intestinal morphology, increases the villi number and length as well as prevents apoptosis. As DPP-IV inhibitor, PTY 2 acts as an effective herbal agents for the treatment of intestinal diseases (Figure 5).

**Fig. 5.**
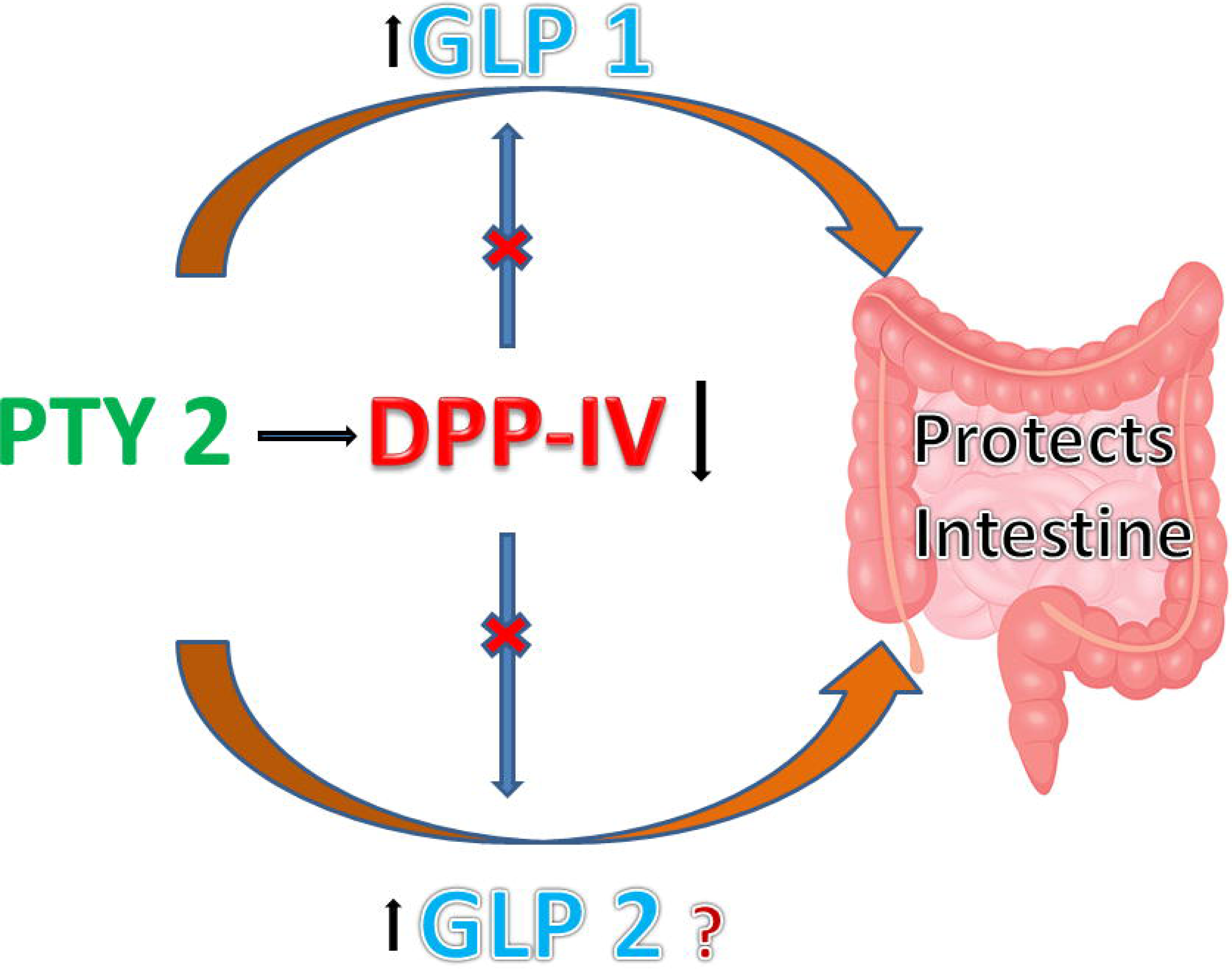
Signaling pathway of PTY-2 acting against diabetes induced intestinal damage.

## 6. CONSENT FOR PUBLICATION

Not Applicable

## 7. AVAILABILITY OF DATA AND MATERIALS

The data and materials supporting the conclusions of this work are included in the article.

## 9. ABBREVIATIONS

PTY 2 (*Pueraria tuberosa* water extract), DPP-IV (Dipeptidyl Peptidase IV), STZ (Streptozotocin), SOD (*Superoxide dismutase*).

## 10. FUNDING

This is a project of the Department of Biotechnology (DBT) (P-07/547) and University Grant Commission (UGC-RGNFD) (F./2013-14/RGNF-2013-14D-GEN-UTT-50872), Government of India.

### 11. ACKNOWLEDGMENT

We are thankful to Prof. Shail Kumar Chaube and his students, Department of Zoology, Institute of Sciences, BHU, for their help in tunnel assay.

## 12. ETHICS APPROVAL AND CONSENT TO PARTICIPATE

Institute Ethical Committee (Dean/2015/CAEC/1266), Institute of Medical Sciences, Banaras Hindu University has approved the overall protocol.

